# Selection efficacy differs between lifestages in the haploid-diploid *Marchantia polymorpha* subsp. *ruderalis*

**DOI:** 10.1101/2024.09.06.611587

**Authors:** Tianyu Shen, Philippe Gadient, Justin Goodrich, Hannes Becher

**Affiliations:** Morgan State University, Baltimore, USA; Institute of Molecular Plant Sciences, Schools of Biological Sciences, University of Edinburgh, Max Born Crescent, Edinburgh EH9 3BF, UK; The Roslin Institute, Royal (Dick) School of Veterinary Science, University of Edinburgh, Midlothian EH25 9RG, UK

## Abstract

Ploidy has profound effects on evolution. Perhaps the most compelling effect is related to dominance: In cells with more than one genome copy, the effects of non-dominant deleterious variants are ‘masked’ to some extent, leading to a reduction in the efficacy of selection. This, in turn, leads to increased levels of nucleotide diversity, and it may also affect nucleotide substitution rates.

To test this predicted association between ploidy and the efficacy of selection, we studied genome scale per-gene patterns of genetic diversity and divergence in the genome of the haploid-diploid bryophyte Marchantia. We treated lifestage gene expression bias as a continuous covariate and accounted for genomic autocorrelation patterns using smoothing splines in a general additive regression model (GAM) framework.

Consistent with a lower efficacy of purifying selection, we found increased levels of sequence diversity, Watterson’s *θ*, and net divergence at non-degenerate sites in genes with diploid-biased gene expression. These genes also showed reduced levels of codon usage bias. In addition, we found chromosome 5 to be an outlier with overall decreased levels of diversity, the site-frequency spectrum skewed towards common alleles, and increased linkage disequilibrium.

In this study, we show the utility of generalized additive models in population genomics, and we present evidence for a ploidy associated difference in the efficacy of selection. We discuss parallels to the evolution of (diploid) sex chromosomes and why the patterns observed are unlikely to be mediated by masking.

## 1 Introduction

Ploidy has profound effects on the pace, course, and constraints on evolution in populations and species (Fox et al., 2020; Pandit et al., 2011; Schmickl and Yant, 2021). It varies at a fine scale across the tree of life, and many species comprise individuals of different ploidy levels (Kolář et al., 2017). 16 % of all flowering plants contain multiple ploidy levels (Rice et al., 2015). This includes crops like potato (De Jong and Tai, 1977) and sweet potato (**pereira_sweetpotato_2023**) as well as invasives/colonisers such as common reed (Pyšek et al., 2020), cordgrass (Fortune et al., 2008; Renny-Byfield et al., 2010), and monkeyflower (Salony et al., 2024). Ploidy can thus be seen as an individual-level property. In addition, ploidy commonly changes during a species’ lifecycle as all sexual species, by definition, go though lifecycle stages of different ploidy levels. In plants, these lifestages are called ‘gametophyte’ and ‘sporophyte’, with the sporophyte having twice the ploidy level of the gametophyte. In the simplest case, the gametophyte is haploid and the sporophyte diploid.

When there are multiple genome copies, the concept of dominance becomes relevant. Any allele that is not fully dominant, i.e. with a dominance coefficient *h* ≤ 1, should then be ‘masked’ to some extent if at least another ‘wild-type’ allele is present at the same locus. Increased ploidy should thus coincide with a reduced efficacy of selection and increased genetic load of deleterious mutations (Kondrashov and Crow, 1991). A reduced efficacy of selection should lead to higher levels of nucleotide diversity in any one population, due to an increased number of segregating slightly deleterious variants. A reduced selection efficacy can also affect non-synonymous divergence, *d*_*A*_, for instance by reducing the number of advantageous substitutions relative to neutral ones. This is the theoretical explanation of faster-X effect (Charlesworth, Coyne, et al., 1987), where the X chromosome, whose loci are effectively haploid in males of many species, shows higher non-synonymous substitution rates than autosomes.

Carrying out within-genome comparisons is a powerful approach for studying the effects of ploidy on the efficacy of selection because it cuts out the potential for confounding with species or population idiosyncrasies. Much information has been gained from studies comparing genes on sex chromosomes to those located on autosomes (e.g. Hu et al., 2013; Langley et al., 2012; Thornton and Long, 2002; Zhou and Bachtrog, 2012) and by comparing X-chromosomal genes with differential expression sex bias (e.g. Campos, Johnston, et al., 2018; Rousselle et al., 2016). But sex chromosomal genes evolve under their own specific conditions and constraints and they only account for a limited fraction of the genome (Avery, 1984). A much larger proportion of the genome can be analysed in haploid-diploids. These are organisms that have separate haploid and diploid lifestages, both showing mitotic growth and developmental differentiation, accompanied by differential gene expression. The haploid-diploid setting allows one to test whether lifestage differences in gene expression, and thus ploidy, significantly affects nucleotide diversity and divergence. Accounting for a significant fraction of the tree of life, haploid-diploids cover a wide spectrum as to the relative importance and persistence of the two lifestages (Bourdareau et al., 2021; Coelho et al., 2007).

One possible issue with within-genome comparisons is that there tend to be autocorrelation patterns: Close-by genes tend to be similar in aspects such as their nucleotide diversity or the skew of their site-frequency spectrum. This observation has several potential causes which are not mutually exclusive, including selection at linked sites (Charlesworth, Morgan, et al., 1993; Maynard Smith and Haigh, 1974), which is mediated by the local rate of recombination and the strength of which may thus change over time (Booker et al., 2022). Further cause include genome-wide variation in mutation rates (Barroso and Dutheil, 2023; Castellano et al., 2020; Hodgkinson et al., 2009; López-Cortegano et al., 2021; Smith et al., 2018), and the mutagenic potential of recombination (Halldorsson et al., 2019; Hinch et al., 2023). Genomic autocorrelation must be taken into account if reliable conclusions are to be drawn from within-genome comparisons.

To investigate whether lower ploidy is associated with a higher efficacy of selection (in analogy the the faster-X effect), we chose to study the haploid-diploid liverwort *Marchantia polymorpha* subsp. *ruderalis*. This taxon has long served as a scientific model organism (Bowman, 2016). It is commonly known, and referred to below, as Marchantia. Marchantia is a member of the bryophytes, the non-vascular land plants. Typical for a bryophyte, Marchantia’s haploid gametophyte lifestage dominates the lifecycle while its diploid sporophyte stage is always dependent on the gametophyte, persisting for a few weeks and eventually releasing large quantities of meiotic spores. See Bowman et al. (2022) for details on Marchantia’s lifecycle. Marchantia is a cosmopolitan ruderal taxon, which disperses via airborne meiotic spores and vegetative gametophytic propagules called ‘gemmae’. Both are haploid. Surprisingly, and despite the effective ploidy differences experienced by genes with differential lifestage expression differences, previous genome scale studies in bryophytes did not find significant effects of lifestage expression differences on the efficacy of selection (Szövényi et al., 2013) or the rate of molecular evolution (Linde et al., 2021).

The purpose of this study is twofold: (1) we follow up on the question of whether there are ploidy effects on the efficacy of selection in bryophytes using novel data and methodology, and (2) we demonstrate the utility of the generalized additive model (GAM) in population genomics. To achieve this, we analysed genome scale patterns of genetic diversity in the bryophyte Marchantia. We characterised nucleotide diversity, linkage disequilibrium, codon usage bias, and the net divergence from another subspecies called *Marchantia polymorpha* subsp. *polymorpha*. We then characterised the distribution of per-gene lifestage expression differences, and assessed how it was associated with diversity and divergence patterns. We found that levels of nucleotide diversity and divergence at non-synonymous sites were significantly increased in genes whose expression was biased towards the diploid sporophyte lifestage. We discuss possible reasons for this observation and parallels to the evolution in diploid sex chromosome systems.

## 2 Results

### 2.1 Data overview

We generated 38.3 Gbp of short-read whole-genome resequencing data (2.0-3.4 Gbp per individual) from 13 individuals of *Marchantia polymorpha* subsp. *ruderalis* (Marchantia). These individuals had been collected in southern Edinburgh, UK, with the sampling area spanning approximately 10 km, see Figure 1. We also reanalysed 115 Gbp of data generated by Sandler et al. (2023). These were from Canadian samples of both Marchantia (15 individuals) and the subspecies *M. p*. subsp. *polymorpha* (8 individuals). The Canadian range spanned a much larger area of several 100 km across. The proportion of successfully mapped reads was considerably higher for our data, as expected for DNA extracted from axenically cultivated samples. The mapping rates of our data ranged from 71 to 96 % with a mean of 93 %, resulting in comparable overall mapping depths for both datasets. After k-mer based masking of repeat regions in the reference assembly, 143 Mbp of autosomal regions (68.6 %) remained for us to carry out variant calling. Screening unmasked coding regions, we identified 3 442 987 fully degenerate (4-fold) and 12 471 802 non-degenerate (0-fold) nucleotide sites.

**Figure 1:**
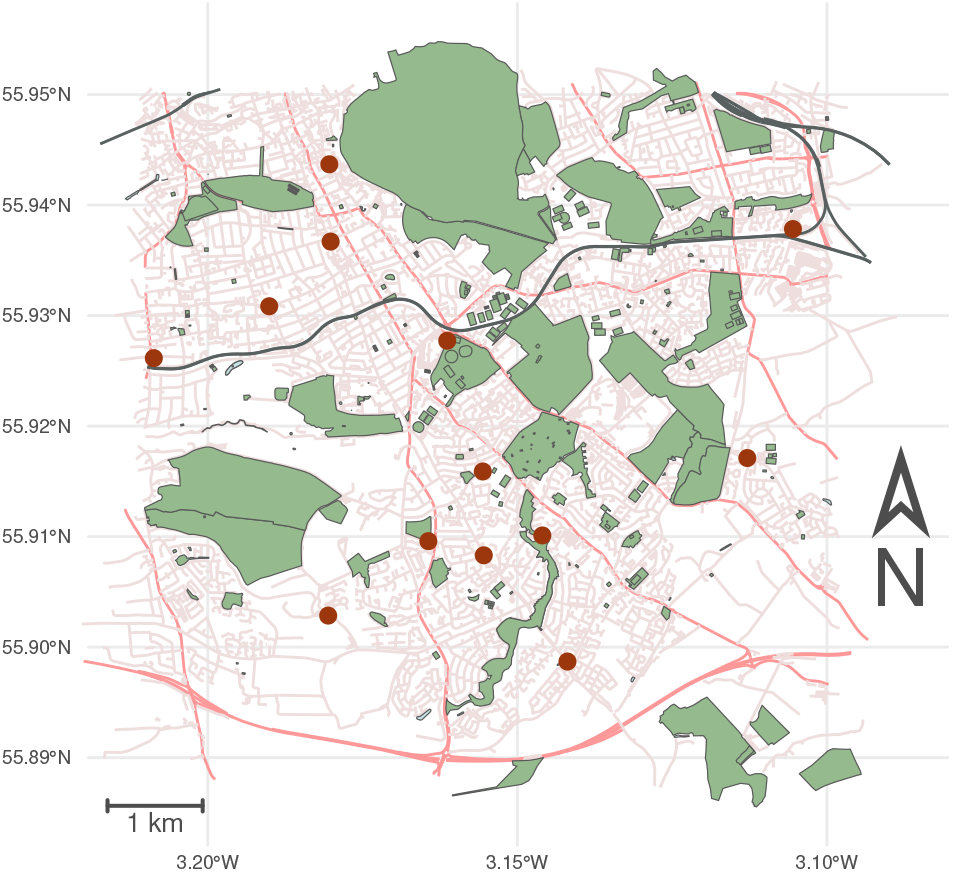
Sampling sites of Marchantia (maroon dots) in southern Edinburgh, UK.

Despite Marchantia’s capacity for clonal reproduction and the modest size of our sampling area, we found that all individuals were genetically distinct. Pairwise per-nucleotide differences at 4-fold sites ranged from 0.001 76 to 0.002 55, suggesting an appreciable degree of sexual reproduction. This is in line with the frequent observation of Marchantia’s iconic reproductive structures, the gametangiophores.

### 2.2 Marked differences in chromosomal diversity patterns

Summary statistics of genetic diversity (Table 1) were in line with the numbers reported by Sandler et al. (2023). However, we computed these on a per-chromosome basis, which revealed considerable inter-chromosomal variation. Chromosome 5 turned out to be an outlier with reduced levels of both genetic diversity, measured as nucleotide diversity, *π*, and the standardised number of segregating sites, *θ*_*w*_. Chromosome 5 also showed an increased skew of the site-frequency spectrum towards common alleles indicated by reduced values of Δ*θ*_*w*_ = 1 − *π/θ*_*w*_ (Becher et al., 2020), see Figure 2A-C. The same chromosome also showed increased levels of linkage disequilibrium, see Figure 2D.

**Table 1:** Per-chromosome summary statistics for Marchantia from Edinburgh. All statistics were computed per gene and then averaged per chromosome. Each is given for both fully degenerate (subscript 4) and non-degenerate sites (subscript 0), with three significant digits. *π* – average number of pairwise nucleotide differences; *θ*_*w*_ – standardised number of segregating sites; Δ*θ* = 1−*π/θ*_*w*_ – a measure of the skew of the site-frequency spectrum; *d*_*A*_ – net divergence from subspecies to *M. p. polymorpha, n* – number of genes considered.

| Chr | $\pi_0$ | $\pi_4$ | $\theta_{w0}$ | $\theta_{w4}$ | $\Delta\theta_{w0}$ | $\Delta\theta_{w4}$ | $d_{A0}$ | $d_{A4}$ | $n$ |
| --- | --- | --- | --- | --- | --- | --- | --- | --- | --- |
| 1 | 0.000711 | 0.00242 | 0.000731 | 0.00248 | 0.0834 | 0.0176 | 0.00518 | 0.0218 | 1858 |
| 2 | 0.00102 | 0.00269 | 0.00101 | 0.00279 | 0.0694 | 0.0618 | 0.00635 | 0.0236 | 1547 |
| 3 | 0.0011 | 0.00315 | 0.00109 | 0.00316 | 0.0742 | 0.0262 | 0.00546 | 0.0231 | 1515 |
| 4 | 0.000939 | 0.00254 | 0.000976 | 0.00261 | 0.109 | 0.0569 | 0.00651 | 0.0234 | 1427 |
| 5 | 0.000814 | 0.00215 | 0.000795 | 0.00201 | -0.0125 | -0.0507 | 0.00703 | 0.0258 | 1396 |
| 6 | 0.00143 | 0.00368 | 0.00139 | 0.00368 | 0.0543 | 0.0462 | 0.00547 | 0.0188 | 1290 |
| 7 | 0.000909 | 0.00234 | 0.000925 | 0.00239 | 0.0926 | 0.0452 | 0.00659 | 0.0247 | 1172 |
| 8 | 0.000932 | 0.00266 | 0.000954 | 0.00264 | 0.0768 | 0.0434 | 0.00558 | 0.0217 | 1075 |
| all | 0.000971 | 0.0027 | 0.000973 | 0.00272 | 0.0694 | 0.032 | 0.00599 | 0.0229 | 11280 |

**Figure 2:**
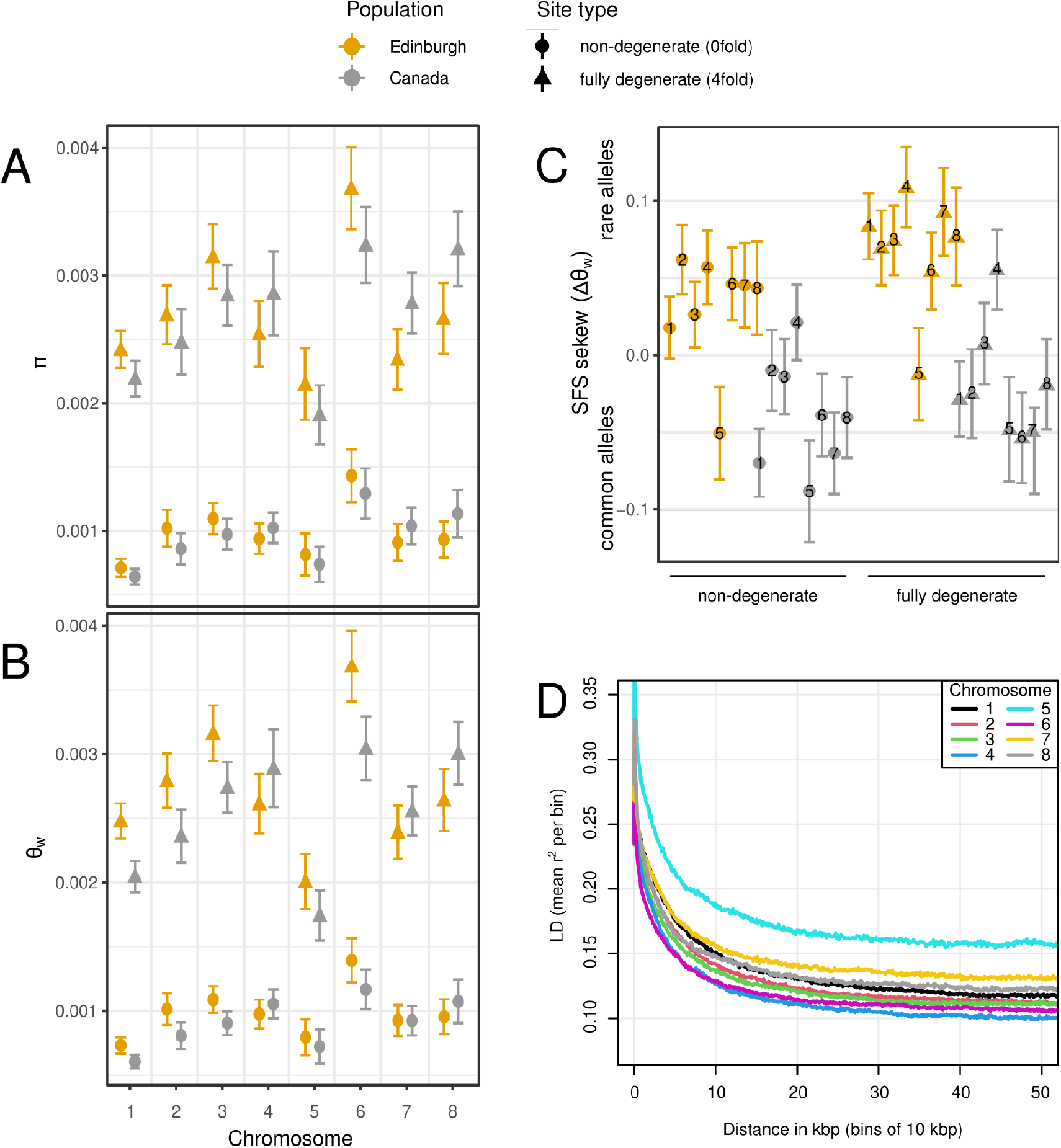
Summary statistics computed per gene, aggregated per chromosome of Marchantia from Edinburgh (mustard) and Canada (grey). Circles indicate the values for non-degenerate, and triangles the values for fully degenerate sites. The error bars indicate 95%-confidence intervals. Panel A shows nucleotide diversity, panel B shows Watterson’s *θ* (the normalised number of segregating sits). A and B share the same x-axis, and the statistics are given per nucleotide. Panel C shows the skew of the site-frequency spectrum as Δ*θ*_*w*_ with higher values indicating an excess of rare alleles. Panel D shows the decay of the linkage disequilibrium measured as the squared correlation coefficient between allele biallelic sites.

There was no significant association between chromosome length and any measure of genetic diversity.

### 2.3 Lifestage expression differences

We chose to measure the per-gene level of lifestage expression bias as the proportion of haploid expression (PHE). This statistic indicates, per gene, what proportion of the overall expression came from haploid tissues. PHE ranges, per definition, from 0 (exclusively sporophytic/diploid) to 1 (exclusively gametophytic/haploid). We found the distribution of the PHE values was shifted towards expression in the gametophyte, which is the predominant lifestage, see Figure 3. A majority of 63 % of genes showed haploid biased expression, i.e. with PHE *>* 0.5. The top 10 % PHE genes were expressed at least 16.5 times more in haploid tissues, whereas the bottom 10 % PHE genes were expressed only at least 3 times more in diploid tissues (arrows in Figure 3). There were 485 genes that were expressed only in haploid tissues, but not a single gene was expressed exclusively in diploid tissues.

**Figure 3:**
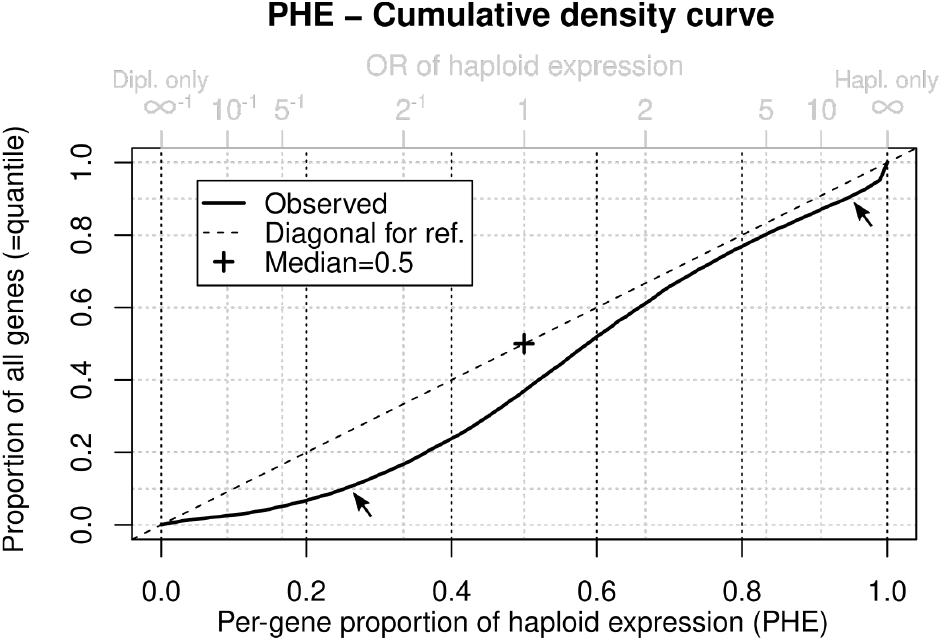
The distribution of per-gene lifestage expression bias, measured as the proportion of haploid expression (PHE). The fat black line shows the cumulative density curve of PHE, ranging from 0 to 1 on the black primary x-axis. The y-axis shows the PHE quantiles. A distribution with a median of PHE = 1/2 (odds ratio=1) would pass through the cross. The dashed diagonal line is given for reference. The arrows indicate the 10^th^ and 90^th^ percentiles, highlighting the distribution’s asymmetry. The grey secondary x-axis applies to the same data, showing odds ratios instead of proportions.

The genomic distribution of PHE was non-random with significant differences between chromosomes (ANOVA, *F*_14067,7_ = 8.87, *p* = 6.08 *×* 10^−11^). While statistically significant, this only accounted for a minute proportion of the variance in PHE. The range of per-chromosome mean PHE values was 0.012, and the adjusted *r*^2^ values were 0.004 when accounting for chromosome effects. It increased to 0.01 when additionally fitting splines to allow for variation in PHE along each chromosome.

### 2.4 Associations between measures of genetic diversity and lifestage expression bias

We then tested whether patterns of genetic diversity and divergence covaried with lifestage expression bias measured as PHE by using generalized additive regression models (GAMs; Wood, 2017). GAMs allow one to combine the usual linear combination of predictor variables used in other regression models with smoothing splines. This is extremely useful, for instance to take account of autocorrelation patterns, which would otherwise violate the regression assumption of independent residuals. In our models, we included covariates of genetic diversity identified in previous studies: overall expression level, tissue specificity, local gene density (Slotte et al., 2011; Wright et al., 2004), number of exons, CDS length, and GC content (Larracuente et al., 2008; Yang and Gaut, 2011). In addition, we included smoothing splines (Wood, 2003) to take account of any autocorrelation of the respective regression outcome variable along the chromosomes.

Below, we report the implied differences between hypothetical haploid and diploid-specific genes which are identical for all other covariates. This is one way of interpreting the coefficients estimates obtained from the regression models. It is important to note that these values differ from the observed values of actual haploid or diploid-specific genes, of which there are few, and which differ not only in their degree of lifestage expression bias but also with respect to all other covariates. In the remainder of the results section, we will focus on the Edinburgh dataset and mention the results obtained from the Canadian dataset for comparison.

#### 2.4.1 Genetic diversity is reduced in gametophyte-biased genes

We found that the pairwise nucleotide diversity at non-degenerate sites, *π*_0_, significantly co-varied with lifestage expression bias (*p* = 3.61 *×* 10^−5^), with higher levels of gametophyte expression bias associated with with lower values of diversity. The coefficient estimate of −0.0907 on log10 scale implies that, all else being equal, the expected level of the nucleotide diversity at non-degenerate sites in a gametophyte specific gene is only 81.1 % of that seen in a sporophyte specific gene (95 % confidence interval: 73.5 %; 89.6 %).

All other covariates, except gene density, also had significant associations with *π*_0_, see Figure 4 (top row, top-left triangles). Tissue specificity of expression, *τ*, was associated with increased diversity, whereas overall expression, number of exons, CDS length, and GC content all were associated with reduced diversity. In analogous models, nucleotide diversity at fully degenerate sites, *π*_4_, was not significantly associated with PHE, but all other covariates were. We observed qualitatively similar patterns when analysing the Canadian dataset, where *π*_0_ was significantly associated with lifestage expression bias (*p* = 0.00218).

**Figure 4:**
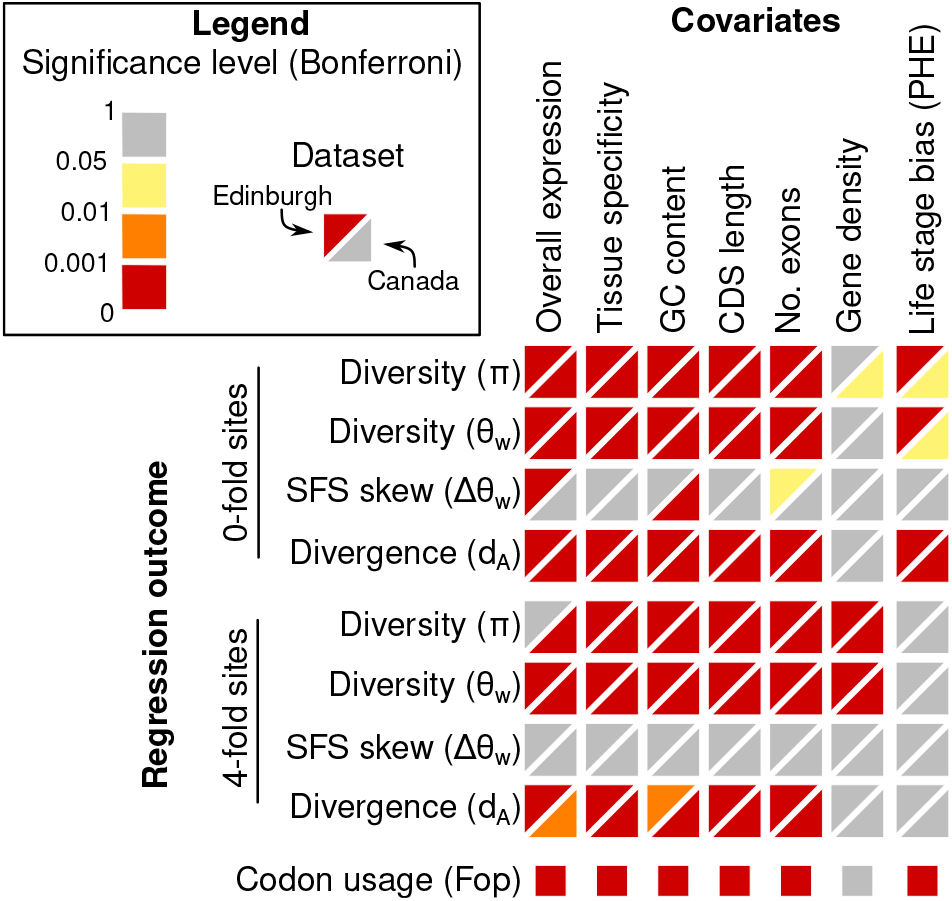
Significance levels of covariates regressed on summary statistics of genetic diversity and divergence for two Marchantia datasets using GAMs. Top-left triangles indicate the Edinburgh and bottom-right the Canadian dataset. Colours represent significance levels with thresholds indicated in the legend. The figure summarises the results of 17 regression models, and the p-values were corrected accordingly (Bonferroni). The significance level of the effect of life stage expression bias, the assessment of which was a key objective of this study, is shown on the very right. Fop (bottom row), the frequency of the optimal codon, which is a measure of codon usage bias, was estimated for the reference assembly. There are thus no separate results for Edinburgh and Canada.

An alternative measure of genetic diversity, *θ*_*w*_, showed similar behaviour to *π*. Increased gametophyte expression bias was significantly associated with lower values of *θ*_*w*0_ (*p* = 2.06 *×* 10^−5^). The coefficient estimate of −0.08 implies that haploid-specific genes have only 83.4 % of the diversity observed in diploid-specific ones (95 % CI: 76.8 %; 90.7 %). All other covariates except for gene density were significantly associated, with all coefficient estimates showing the same sign as for *π*. Expression bias was not significantly associated with the standardised number of segregating fully degenerate sites, *θ*_*w*4_, but all other covariates were. The Canadian dataset showed analogous significance levels.

#### 2.4.2 Codon usage is more optimal in haploid biased genes

The per-gene frequency of the optimal codon (Fop) was significantly increased in haploid biased genes (*p* = 1.28 *×* 10^−20^). The implied difference in Fop between haploid and diploid specific genes was 0.0241.

#### 2.4.3 The site-frequency spectrum is not skewed by lifestage expression bias

We did not find significant associations between lifestage expression bias and the skew of the site frequency spectrum, measured as Δ*θ*_*w*_, neither for fully degenerate nor non-degenerate sites. This was so for both the Edinburgh and Canadian sample. Overall, the site-frequency spectra of the Edinburgh samples tended to be skewed towards rare alleles, showing positive Δ*θ*_*w*_, whereas the Canadian sample tended to show skew towards common alleles, negative Δ*θ*_*w*_, see Figure 2C.

#### 2.4.4 Divergence at non-synonymous sites is reduced in gametophyte-biased genes

Using population data, we were able to estimate the net divergence, *d*_*A*_, between Marchantia and *M. p*. subsp. *polymorpha*. The net divergence at non-degenerate sites was significantly reduced in genes with higher lifestage expression bias. The coefficient estimate of −0.130 on log10 scale implies a reduction of the net divergence at non-degenerate sites in haploid-specific genes to 74.2 % of the level of diploid-specific genes (95 % CI: 67.7 %; 81.3 %). All other covariates were significant, see Figure 4, with the same direction of effects as we found for *π*. Net divergence at fully degenerate sites did not covary with lifestage expression bias while all other associations remained significant. The picture for the Canadian samples was analogous.

## 3 Discussion

In this study, we characterised patterns of genetic diversity in a sample of Marchantia from Edinburgh, UK, to test whether there is an association between genes’ lifestage expression bias and their patterns of genetic diversity and divergence. We also reanalysed a sample of Marchantia from Canada (Sandler et al., 2023). We found that genes with expression bias towards the haploid gametophyte lifestage showed, at non-degenerate sites, significantly reduced levels of genetic diversity and sequence divergence. Below, we first point out methodological differences from previous studies. We then discuss our findings in light of the expected efficacy of selection and ploidy.

### 3.1 Methodological advances

A higher efficacy of purifying selection at the haploid gametophyte lifestage has been reported in several systems (Arunkumar et al., 2013; Gossmann et al., 2014; Lipinska et al., 2019; Tanja Slotte et al., 2010). A higher efficacy of purifying selection on the X chromosome, which is haploid in males, is one explanation for the commonly observed faster X effect, which we mentioned in the introduction. In closer analogy with the haploid-diploid setting, the relative rate of adaptive substitutions, *ω*_*α*_, on the X chromosome is highest in male-biased genes in Drosophila, which are effectively haploid when expressed (Campos, Johnston, et al., 2018). However, two previous studies on bryophytes paint a different picture. Linde et al. (2021) concluded that there was no evidence for lifsestage differences in selection pressure as there was no increased *d*_*N*_ */d*_*S*_ in the haploid lifestage in Marchantia. Studying the moss *Funaria hygrometrica*, Szövényi et al. (2013) found a complex relationship between *d*_*N*_ */d*_*S*_ and lifestage expression bias – genes specific to either the sporophyte or gametophyte showed lower *d*_*N*_ */d*_*S*_ values than unspecific genes. The authors explained this pattern by also taking into account tissue specificity, which tends to be lower in genes with lifestage specific expression.

Our study here differs in three key aspects. First, we used a continuous measure of lifestage expression bias, which we call the proportion of haploid expression. We chose this approach because the alternative – classifying genes into expression bins – always comes with a degree of arbitrariness and because classification does not allow for measurement error (noise). The more tissue samples are analysed in an experiment, the lower becomes the probability that a gene will be counted as lifestage specific. In addition, if there are bins for lifestage specific expression, the probability of a gene’s assignment to these bins will also depend on the relative number of tissues sampled for each lifestage. In the expression dataset that we used (Marpolbase Expression, https://mbex.marchantia.info/), there were far more gametophyte samples than sporophytic ones, making it more likely for low-expressed genes to appear gametophyte-specific by random chance. Indeed, we observed hundreds of gametophyte-specific genes but none that was sporophyte specific. This pattern is of cause plausible given the biology of Marchantia, where the gametophyte lifestage is small and has a short duration. But the pattern is likely exacerbated by the tissue sampling available in the expression database. By treating expression bias as continuous, we treat it as data, which comes with uncertainty. Any gene with observed lifestage specific expression is not categorically different from any other genes. Lifestage specific genes just sit at the extremes of the continuous PHE scale, which ranges from 0 or 1.

Second, we generated population data, which allowed us to analyse patterns of nucleotide diversity within Marchantia. Population data are also required to estimate *d*_*A*_, the net divergence (Nei, 1987; Nei and W.-H. Li, 1979). In the absence of population data, it is possible to estimate the pairwise nucleotide divergence, *d*_*xy*_, from a pair of genome or transcriptome sequences. But the value of *d*_*xy*_ is affected by both the divergence between and diversity within the populations compared, even when comparing only two sequences. The distinction between *d*_*xy*_ and *d*_*A*_ is thus more important when comparing recently-diverged taxa where much of the expectedly low *d*_*xy*_ estimate is due to within-population (possibly ancestral) diversity.

Third and finally, we took account of ‘spatial’ autocorrelation along chromosomes by fitting smoothing splines using general additive models (GAMs). Ignoring autocorrelation leads to non-independent residuals in regression analyses, which can compromise significance tests and lead to spurious conclusions. Early studies with small numbers of loci could safely ignore the possibility of autocorrelation (Begun and Aquadro, 1992; Payseur and Nachman, 2002) without detrimental effect. More recent genome-wide studies commonly use window-based approaches (Slotte et al., 2011), take the ratio of the same statistic at non-degenerate and fully degenerate sites such as *π*_0_*/π*_4_, or explicitly take into account likely causes of autocorrelation (Becher et al., 2020; Campos, Halligan, et al., 2014). There are several advantages of fitting smoothing splines within the GAM framework. Being a non-parametric approach, it is not necessary to make assumptions as to the cause of autocorrelation. Also, because GAMs are ‘fully-fledged’ regression models, they are more flexible than rank regression. GAMs allow for more complex hypothesis testing and the inclusion of otherwise possibly confounding covariates. Using spline-based smoothing on chromosomal positions is arguably less arbitrary than selecting a window size for window-based analyses. Accounting for autocorrelation by smoothing, in turn, allows for the separate analysis of statistics computed on non-degenerate and fully degenerate sites rather than their ratios.

### 3.2 The efficacy of selection

It may appear plain that the efficacy of selection should differ between genes with different degrees of life cycle expression bias. The effect of masking in diploids or ‘unmasking’ in haploids (Beaudry et al., 2020) and the associated higher efficacy of selection in haploids is an extremely compelling argument in favour of such differences. There is however another mechanism potentially affecting the efficacy of selection in the opposite way. Because there is no additional genome where expression might happen at the same locus, haploids may show a higher degree of variation in gene expression levels (‘expression noise’) than organisms of higher ploidy (Cook et al., 1998). Wang and J. Zhang (2011) have compared the consequences of this phenomenon, which has been observed in yeast, to a depression of the effective population size in haploids relative to diploids. Increased expression noise in haploids would reduce the efficacy of selection. Expression noise might perhaps be mitigated by endopolyploidy, a condition where multiple genome copies are present in somatic cells. Endopolyploidy is common in plants and mosses in particular (Bainard, Newmaster, and Budke, 2020) but not known in the liverworts (Bainard and Newmaster, 2010) and is thus unlikely to matter for Marchantia. However, while important in yeast, is it not clear how relevant expression noise is in multicellular organisms Wang and J. Zhang (2011). For instance, gene expression from the functionally haploid mammal X chromosomes does not seem to be more noisy than from autosomes (Yin et al., 2009).

Because directional selection reduces the frequency of deleterious alleles, an increased efficacy of selection should lead to reduced nucleotide diversity. We observed the expected pattern at non-degenerate sites as *π*_0_ was significantly reduced in genes with haploid-biased expression. Selection effects at fully degenerate sites are generally much weaker, leading to overall higher values of *π*_4_ than *π*_0_, a pattern observed by Sandler et al. (2023), which we confirmed here. Fully degenerate sites may be affected by weak direct selection or by section acting on linked nearby sites, the strength of which depends on the local, and possibly historic, recombination rate (Booker et al., 2022; Campos, Halligan, et al., 2014). Both effects should be affected by any difference in the efficacy of selection and there could in principle be a depression of *π*_4_ at haploid-biased genes. We did not observe such a pattern. We did however find other evidence for a difference in the efficacy of direct selection on degenerate sites. Codon usage differed, with the frequency of the optimal codon significantly higher at haploid-biased genes. This is in analogy to findings in Drosophila, where the effective codon usage bias is larger on the X than on the autosomes (Campos, Zeng, et al., 2013). Overall, these observations are evidence for a higher efficacy of selection at the gametophyte lifestage in Marchantia.

The efficacy of selection may also affect nucleotide substitution rates. But this effect is expected to be complex because nucleotide divergence may be due to adaptive, neutral, or even (slightly) deleterious substitutions. Depending on the distribution of fitness effects of mutations, a higher efficacy of selection might simultaneously reduce the number of slightly deleterious substitutions and increase the number of adaptive substitutions that overcome the effect of random genetic drift. For this reason *d*_*N*_ */d*_*S*_, sometimes called the ‘rate of evolution’, has been observed to be reduced in species with an increased efficacy of selection (Galtier, 2016).

We observed a significant decrease of non-synonymous substitutions between Marchantia and *M. polymporpha* subsp. *polymorpha* in genes with haploid expression bias. Considering that the proportion of adaptive mutations among all non-synonymous substitutions, *α*, was estimated to be small and not significantly different from zero (Sandler et al., 2023), it would seem plausible to attribute this pattern to a decreased number of slightly deleterious substitutions in genes with gametophyte-biased expression. This is analogous to an observation made in some butterflies. In butterflies, females are the heterogametic sex, carrying only one Z chromosome. Z-chromosomal genes with female-biased expression are thus functionally haploid just like autosomal genes with strong gametophyte expression bias in Marchantia. Rousselle et al. (2016) found that the rate of non-adaptive substitutions in Z-chromosomal genes with female expression bias was reduced compared to male biased ones, leading to an overall reduction of the rate of non-synonymous substitutions in haploid biased genes as we observed in Marchanita. While we found evidence for an increase in the efficacy of selection in haploid biased genes, this is not entirely analogous to the faster-X effect.

While the patterns we report suggest an increased efficacy of selection at the haploid gametophyte lifestage, it is important to note that our study cannot assess whether there is as causal relationship between ploidy and the efficacy of selection.

### 3.3 Within genome comparisons and the effects of ploidy

One rationale for studying the effect of lifestage expression bias is to test the effect of ploidy while excluding potentially confounding effects stemming from species or population idiosyncrasies, as we mentioned in the introduction. In the same organism, all autosomes must go through the same life cycle stages, experiencing the same changes in ploidy and the same levels of genetic drift. Any systematic differences between genes with expression bias towards one stage would thus necessarily be correlated with with ploidy.

Besides ploidy, the basis for proposing effects of unmasking and expression bias, what else differs between the lifestages? One difference is the relative length of, and the number of cell divisions in, each stage. If some gene’s expression were biased towards the shorter life cycle stage, that gene would experience less transcription coupled repair. Transcription coupled repair is a repair mechanism that removes DNA adducts, which would otherwise cause mutations during DNA replication (Hanawalt and Spivak, 2008; Nicholson et al., 2024). As a consequence, genes that are mainly active in the shorter life cycle stage may experience higher mutation rates. In bryophytes, the shorter lifestage is the sporophyte. The effect of reduced transcription coupled repair should thus act to increase levels of nucleotide diversity in the diploid lifestage, similar to masking. An important difference from masking is that lack of transcription coupled repair should also increase levels of synonymous diversity, which we did not observe.

A second aspect differing between the lifestages may be their population sizes. In bryophytes, one haploid gametophyte can, and usually does, carry numerous sporophytes. The realised lifestage specific population size will depend on numerous factors such as availability of suitable gametes, nutritional status of the carrying gametophyte, and (in dioicous species) the sex ratio. If there is a systematic difference in lifestage specific population size, that should affect the efficacy of selection of genes with biased expression (Bessho and Otto, 2017). Again, this should also affect levels of nucelotide diversity of fully degenerate sites, which we did not find to differ.

Third, even if genes are expressed in and under selection at both lifestages, this selection may be of antagonistic nature. Such ploidally antagonistic selection, may also interact with sexually antagonistic selection, can lead to increased diversity in some genes without lifestage expression bias (Immler et al., 2012). This could in principle cause a non-monotonic association between lifestage expression bias and nucleotide diversity.

Fourth, in many haploid-diploid species there is the possibility for life cycle short cuts, with the extreme being the emergence of asexuality (Hoshino et al., 2024). Such short cuts are common in bryophytes where there is vegetative propagation in the gametophyte stage. If the balance between vegetative and sexual reproduction is under genetic control, as modelled by Scott and Rescan (2017), then there are autosomal alleles whose average time spent in each lifestage differs from that of other genes. The respective loci would be expected to show deviant patterns of genetic diversity. However, this would presumably affect only a small number of loci, not skewing the overall picture.

Fifth, recent research has shown that evolutionary rates covary with gene age (Moutinho, Eyre-Walker, et al., 2022) and protein features such as ‘relative solvent accessibility’ and ‘intrinsic disorder’ (Moutinho, Trancoso, et al., 2019). While these are important determinants for the level of divergence between taxa, it seems less likely that these would covary with lifestage expression bias.

Sixth and most importantly, there may be imprinting which causes parent-of-origin bias to allele expression in the sporophyte stage. The potential for imprinting has been pointed out by Haig (2013) and Haig and Wilczek (2006) in relation to life-stage conflict in bryophytes. Recent work has uncovered extreme imprinting in Marchantia, with expression of most sporophyte-expressed genes being strongly biased towards alleles inherited from the maternal gametophyte (Montgomery, Hisanaga, et al., 2022). Such a drastic expression reduction of paternal alleles is bound to reduce the amount of selection acting on these, in particular if the respective gene’s expression shows a strong sporophyte bias. Assuming that deleterious alleles tend to be rare and thus mainly to be present in haploid state, imprinting would approximately halve the selection acting on a sporophyte expressed deleterious allele. Masking would modify the selection by the dominance coefficient *h*. Depending on the value of *h* of mutant alleles in Marchantia, imprinting could have a stronger (*h* ≥ 1*/*2), similar (*h* ≈ 1*/*2), or weaker effects (*h* ≤ 1*/*2) than masking on the reduction of the efficacy of selection. Nevertheless, both imprinting and masking effects should act in the same direction, reducing the efficacy of selection in the sporophyte compared to the gametophyte and thus increasing diversity in the sporophyte compared to the gametophyte.

Because there are numerous phenomena that covary with ploidy, notably the massive imprinting at the sporophyte stage, it is not possible to attribute the observed lifestage differences in selection efficacy to the effects of masking.

### 3.4 Conclusion and outlook

Using novel methodology, we found significant associations between patterns of diversity and divergence on one hand and the degree of per-gene lifestage expression bias on the other, in the haploid-diploid liverwort Marchantia. We see this as evidence for a higher selection efficacy at the gametophyte lifestage of Marchantia. This is effect is unlikely to be caused by the masking of deleterious mutations in the diploid lifestage because of the pervasive imprinting observed in Marchantia’s sporophytes (Montgomery and Berger, 2024).

Recent and ongoing theoretical research keeps adding to our understanding of evolution in haploid-diploids, where allelic fixation probabilities can differ considerably from species with other life cycles (Bessho and Otto, 2017, 2022) and lifestage expression bias may affect the architecture of local adaptation (Zwaenepoel et al., 2024) in analogy to sex chromosomal expression (Fraïsse and Sachdeva, 2021; Lasne et al., 2017). Promising new simulation tools have become available (Haller and Messer, 2023; Sorojsrisom et al., 2022), which may help isolating the effect of ploidy while varying other life cycle parameters. It is an exciting time to be working on haploid-diploids.

## 4 Methods

### 4.1 Plant collection, cultivation, DNA extraction, and sequencing

We collected 13 individuals of *Marchantia polymarpha* subsp. *ruderalis* in southern Edinburgh (UK). We then established axenic lines from single gemmae sterilised with NaDCC (Duckett et al., 2004), which we grew on plates of a modified Johnson’s medium (Mulvey and Dolan, 2023) until the thalli reached approximately 2 cm.

We extracted genomic DNA using a phenol chloroform protocol (Lu et al., 2024). The DNA was sequenced at Novogene (Cambridge, UK) to yield approximately 5X coverage depth (discounting for PCR duplicates).

### 4.2 Genomic sequencing data and related bioinformatic analyses

We downloaded the newly generated sequencing data in FASTQ format from Novogene’s web service. We also downloaded the data generated by Sandler et al. (2023). We trimmed all reads using Skewer (Jiang et al., 2014), discarding all 3’ nucleotides with a PHRED quality score less then 30. We downloaded from https://marchantia.info/ the the MpTak v6.1 reference assembly and annotation, and we aligned all sequencing reads using bwa-mem2 (Vasimuddin et al., 2019). We coordinate-sorted the resulting BAM files using SAMtools (H. Li and Durbin, 2009), marked duplicates using Picard tools (https://broadinstitute.github.io/picard/), and finally called variants with Freebayes (Garrison and Marth, 2012).

To restrict our downstream analyses to reliable single-copy regions, we generated a BED file of repetitive regions using the k-mer based masking program Red (Girgis, 2015). Starting from the reference assembly and annotation, we used Simon Martin’s Python script codingSiteTypes.py from the repository https://github.com/simonhmartin/genomics_general to identify fully degenerate and non-degenerate nucleotide sites. We developed Python code using the package scikit-allel (Miles et al., 2023) to compute population genetic statistics per gene.

To assess the decay of linkage disequilibrium, we used PopLDdecay (C. Zhang et al., 2019). Because this program runs only on diploid VCF files, we generated a phased ‘diploid’ VCF by joining pairs of individuals into diploid variant call columns.

### 4.3 Gene expression statistics

We downloaded the Marpolbase Expression data from https://marchantia.info/mbex/ (accessed 11th April 2023). We then removed samples that might have contained RNA from both sporophyte and gametophyte tissues. We retained only genes with a summed TPM value *>* 10. We then averaged transcription levels per gene and tissue. In a second step, we averaged per-gene tissue means per lifestage. We then divided the average gametophyte expression values by the sum of the average expression values from gametophyte and sporophyte, to obtain the proportion of haploid expression (PHE). PHE being a proportion, its values ranged from 0 to 1. We also computed the overall average expression value for each gene as the average of the per-tissue values. We finally computed the tissue specificity of gene expression, *τ* (Yanai et al., 2005), for each gene. To compute *τ*, we used haploid tissues only to avoid excessive correlation with PHE.

### 4.4 Other per-gene statistics

We computed each of the following statistics for each gene, separately for non-degenerate and fully degenerate sites. We computed nucleotide diversity using the scikit-allel package. We assessed the skew of the site-frequency spectrum both using the statistic Δ*θ* (Becher et al., 2020), which is a simple function of the nucleotide diversity and standardised number of segregating sites Δ*θ* = 1 − *π/θ*_*w*_. We also assessed the net divergence, *d*_*A*_,(Nei, 1987; Nei and W.-H. Li, 1979), between Marchantia and *M. p*. subsp. *polymorpha*. We obtained this by computing *d*_*xy*_ and then subtracting half of each subspecie’s nucleotide diversity. To compute the frequency of the optimal codon, we used codonW (Peden, 1999).

### 4.5 Statistical analyses

We used R (R Core Team, 2023) for all statistical analyses. To analyse the per-gene data and its association with expression in different life stages accounting for background variation in levels of genetic diversity we used the package mgcv (https://CRAN.R-project.org/package=mgcv). In the analyses of nucleotide diversity, *θ*_*w*_, and *d*_*A*_, we used log_10_ transformation on the outcome variable.

## 5 Acknowledgements

HB is indebted to Christelle Fraïsse for discussions that lead to this inception of this study, which begun as TS’s Bioinformatics MSc project at the University of Edinburgh’s School of Biological Sciences. HB is grateful to Alex D Twyford who allowed him to take on two MSc students during his time at the Twyford Lab. We thank Matthew Hartfield, Konrad Lohse, and Brian and Deborah Charlesworth for their comments on an earlier version of this manuscript. This work has made use of the resources provided by the Edinburgh Compute and Data Facility (ECDF) (http://www.ecdf.ed.ac.uk/).

## 6 Data availability

The sequencing reads generated are available from the sequence read archive, BioProject PRJNA1121287. The analysis code is available from Zenodo: www/gggbbb. [Both available upon acceptance of this manuscript.]

## 7 Author contributions

TS carried out bioinformatic analyses and computed per-gene summary statistics. PG and JG supplied access to wetlab infrastructure, helped with plant husbandry, and carried out DNA extractions. HB designed the study and wrote the first draft of the manuscript. All authors read, commented on, and approved the final manuscript.

## 8 Competing interests

The authors declare no competing interests.

## Notes

### Competing Interest Statement

The authors have declared no competing interest.

